# Transcriptome analysis of diverse *Plasmodium falciparum* clinical isolates identifies genes correlating with highly variable expression of merozoite surface protein MSPDBL2

**DOI:** 10.1101/2022.02.18.481051

**Authors:** Suzanne E. Hocking, Lindsay B. Stewart, Sarah J. Tarr, Aline Freville, Kevin K. A. Tetteh, Ambroise D. Ahouidi, Alfred Amambua-Ngwa, Mahamadou Diakite, Gordon A. Awandare, David J. Conway

## Abstract

The merozoite surface protein MSPDBL2 of *Plasmodium falciparum* is under strong balancing selection and is a target of naturally acquired antibodies. Remarkably, MSPDBL2 is expressed in only a minority of mature schizonts of any cultured parasite line, and *mspdbl2* gene transcription increases in response to overexpression of the gametocyte development inducer GDV1, so it is important to understand its natural expression. Here, MSPDBL2 in mature schizonts was analysed in the first *ex vivo* culture cycle of 96 clinical isolates from four populations with varying levels of infection endemicity in different West African countries, by immunofluorescence microscopy with antibodies against a conserved region of the protein. In most isolates, less than 1% of mature schizonts were positive for MSPDBL2 (median of 0.6% overall), but the frequency distribution was highly skewed as nine isolates had more than 3% schizonts positive and one had 73% positive. To investigate whether expression of other gene loci correlated with MSPDBL2 expression, whole transcriptome sequencing was performed on schizont-enriched material from 17 of the clinical isolates with a wide range of proportions of schizonts positive. Transcripts of particular parasite genes were highly significantly positively correlated with MSPDBL2 positivity in schizonts as well as with *mspdbl2* gene transcript levels, with overrepresentation of genes previously implicated as likely to be involved in gametocytogenesis, but not including the gametocytogenesis master regulator *ap2g*. Although MSPDBL2 is apparently not directly involved in sexual commitment, it marks a co-occurring developmental subpopulation that may be functionally distinct within blood stage infections.

## Introduction

The *Plasmodium falciparum* merozoite surface protein MSPDBL2 is one of two MSP3-like proteins with a duffy-binding like (DBL) domain, expressed in schizonts and co-localised with MSP1 on the merozoite cell surface (1, 2). MSPDBL2 is not directly membrane-bound, but appears complexed with MSP1, and can bind to the surface of uninfected erythrocytes (1), suggesting a potential role in invasion. The *mspdbl2* gene occurs at a single locus on chromosome 10 with an intact coding sequence in all *P. falciparum* isolates, although it has multiple stop codons in the related chimpanzee parasite *P. reichenowi* indicating that it is not functional in that species (3, 4), and an orthologue only exists within members of the *Laverania* sub-genus but not in other malaria parasites (5). The gene is highly polymorphic within *P. falciparum,* and analysis of allele frequency distributions in endemic populations indicates that diverse alleles are maintained by strong balancing selection (3, 6, 7), suggesting that it may be a target of immune selection.

MSPDBL2 is a target of naturally acquired antibody responses, and an endemic population cohort study indicates some association with reduced prospective risk of clinical malaria (8), while another study showing purified IgG inhibits parasites in culture (9). However, the vaccine candidacy of MSPDBL2 is uncertain, as it is not only polymorphic but also highly variable in expression. Analysis of schizont-rich cultures of clinical isolates and long-term adapted *P. falciparum* laboratory lines has revealed very low *mspdbl2* transcript levels in most isolates assayed by RT-qPCR (6), or by whole transcriptome analysis (10). Strikingly, MSPDBL2 protein expression is restricted to a small proportion of mature schizonts in each laboratory-adapted parasite line tested by immunofluorescence with specific antibodies (6), although frequencies of MSPDBL2 expression in schizonts of clinical isolates have not been reported. Consistent with being expressed in only a small minority of parasites, experimental disruption of the *mspdbl2* gene does not affect overall asexual parasite growth rates in culture (11).

If MSPDBL2 is restricted to a functionally-important parasite subpopulation, it might still be a target to be considered for vaccination. Interestingly, MSPDBL2 has been shown to bind to the Fc region of IgM, although the function of this is unknown (12). Separately, overexpression of *mspdbl2* has been reported to enhance parasite survival in the presence of some antimalarial drugs in culture (13). Significantly, it has been suggested that *mspdbl2* expression may be correlated with parasite sexual differentiation, as over-expression of the *gdv1* gene in an engineered parasite clone results in marked increase in transcription of *mspdbl2* as well as genes known to be involved in switching to gametocyte development (14). This indicates the importance of studying variation in *mspdbl2* gene expression as well as MSPDBL2 protein expression in schizonts of clinical isolates, and explore whether this is associated with markers of parasite commitment to gametocyte development *in vivo*. It is clear that there is great variation in proportions of parasites committed to gametocyte development among different infections from a single endemic area (15), and transcript markers of ring stage parasites that will develop as gametocytes (16), but apart from the master regulator transcription factor AP2-G (17) there are no confirmed markers of sexually committed schizonts in the previous cycle (18, 19). This study investigates the frequency distribution of expression of MSPDBL2 protein in schizonts of clinical isolates from endemic populations, and tests for correlations with other genes expressed in clinical isolate transcriptomes, including those previously implicated as associated with the gametocytogenesis pathway.

## Materials and Methods

### *P. falciparum* clinical isolates

For analysis of parasite phenotypic and expression variation in natural infections, clinical malaria cases attending local health facilities in Ghana (Kintampo), Mali (Nioro du Sahel), Senegal (Pikine), and The Gambia (Basse) were investigated. The level of malaria infection endemicity varied among these different populations in West Africa, being higher at the sites in Ghana and Mali than the sites in Senegal and The Gambia, which is also reflected in more complex mixed genome *P. falciparum* infections at the more highly endemic sites, as previously shown for these populations (20). Patients aged between 2 and 14 years were eligible if they had uncomplicated clinical malaria, had not taken antimalarial drugs in the 72 hours preceding sample collection and tested positive for *P. falciparum* malaria by lateral flow rapid diagnostic test and slide microscopy. Written informed consent was obtained from parents or legal guardians of participating children and additional assent was received from participating children. Up to 5ml of venous blood was collected in heparinised anti-coagulation BD Vacutainer^®^ tubes (BD Biosciences), and a proportion of the erythrocytes were cryopreserved in glycerolyte and stored at −80°C or in liquid nitrogen before shipment on dry ice to the London School of Hygiene and Tropical Medicine for subsequent culture and laboratory analysis. Ethical approval for the collection and analysis of clinical samples was granted by the Ethics Committees of the Ministry of Health in Senegal, the Ministry of Health in Mali, the Ghana Health Service, the Noguchi Memorial Institute for Medical Research, University of Ghana, Kintampo Health Research Centre, MRC Gambia, and the London School of Hygiene and Tropical Medicine.

### *P. falciparum* schizont preparations from the first cycle of *ex vivo* culture

The clinical blood samples were thawed in batches of eight at a time and introduced into culture *ex vivo,* all isolates being processed in culture in a single laboratory, as described for a previous study of the first *ex vivo* malaria parasite generation from similar clinical samples (21). Giemsa-stained thin blood films were prepared for each isolate initially upon thawing, and later during the second day of culture to assess the developmental progression of parasites into schizogony. Isolates containing schizonts in culture on the second day after thawing were enriched for schizonts by magnetic MACS^®^ separation, and parasites were then allowed to mature in the presence of E64 for 4 hours to prevent schizont rupture, following which cells were harvested by centrifugation, using methods similar to those previously applied to studies of schizonts in continuously cultured parasite lines (10). Erythrocytes containing matured schizonts were prepared for immunofluorescence assays by washing and resuspending in 1% BSA and spotting into individual wells of 12-well slides (Hendley-Essex), air dried and stored with desiccant at −40°C until assay, as performed in previous analysis of schizonts in cultured parasite lines (6).

### Analysis of MSPDBL2 expression in schizonts by immunofluorescence

Staining of schizonts positive for MSPDBL2 by immunofluorescence was performed with the identical method previously used on laboratory-adapted *P. falciparum* isolates, involving incubation with polyclonal mouse serum specific for the conserved N-terminal portion of MSPDBL2, and goat anti-mouse IgG Alexa Fluor^®^ 555 secondary antibody, with Vectashield^®^ mounting fluid containing DAPI to visualise nuclei (6). For each isolate, approximately 1000 mature schizonts (each containing at least 8 nuclei) were counted using DAPI and scored for MSPDBL2 expression using a manual Leica fluorescence microscope with a 100x objective. MSPDBL2 expression was always clearly brightly positive or entirely negative in each individual mature schizont, so that counts of numbers and proportions positive in each preparation were recorded, using same process as described previously (6).

### RNA-seq of schizont-enriched samples of clinical isolates

Parasite schizont-enriched material from individual isolates was stored in either TRIzol^®^ or RNA/ater™ (Thermo Fisher Scientific, MA, USA), and RNA was extracted by phenol-chloroform and cleaned up using the NucleoSpin^®^ RNA XS extraction kit (Macherey-Nagel, Germany). Samples showing successful RNA extraction after checking by Bioanalyzer electrophoresis were reverse transcribed and cDNA amplified using the SmartSeq^®^ v4 Ultra^®^ Low Input RNA Kit for Sequencing (Takara Bio. Inc., Shiga Prefecture, Japan). Successfully amplified samples were prepared for paired end short-read sequencing using the Nextera XT Library Prep kit (Illumina, California, USA), individual libraries being pooled in equimolar amounts at 4nM with up to 12 per pool, and sequencing was performed on an Illumina MiSeq using the 150-cycle MiSeq reagent kit v3. RNA from isolate INV236 which had the highest proportion of MSPDBL2-positive schizonts (73%) was prepared and sequenced as a priority, and following this RNA was extracted from 38 other isolates with varying proportions of MSPDBL2-positive schizonts, of which 25 showed expected cDNA size range profile after reverse transcription and amplification, and 16 of these were selected for sequencing as they had RNA quality RIN score > 6. This yielded a set of 17 isolates with RNA-seq data and matched IFA data on proportions of MSPDBL2-positive schizonts.

Following procedures previously used for RNA-seq analysis of schizont-enriched *P. falciparum* cultures of other isolates (10), whole transcriptome short read sequence data were assembled by alignment mapping to the *P. falciparum* 3D7 version 3.0 reference genome (22) using HISAT2 (23). Gene transcript levels were assessed using the FPKM metric (Fragments Per Kilobase of transcript per Million mapped reads, the number of reads mapping to each gene normalised for the size of the sequencing library and for gene length). Data were analysed using the R package DESeq2 (24), using a masked GFF annotation file that removed the *var, rifin,* and *stevor gene* families from analysis, as described for analysis of previous schizont transcriptome data (10). In addition to exclusion of these three sub-telomeric gene families from analysis, portions of other protein-coding genes that show high allelic diversity (including the highly polymorphic central region of the *mspdbl2* gene) were masked to ensure mapping occurred only in conserved regions of those genes to minimise allelic bias among samples, as previously described for other data (10). Prior to analysis, three independent studies including proteomic or transcriptomic analyses were consulted (14, 25, 26) to identify *P. falciparum* genes considered to be potentially associated with gametocytogenesis, enabling compilation of a list of 119 genes (Supplementary Table S1) used as a set for conducting exploratory correlative analyses of the RNA-seq data within this study. To avoid discovery bias in the present study this needed to be a static list fixed prior to analysis, but is not a reference list for other studies as ongoing research means that any such compilations should be updated and can be subject to different criteria.

### Statistical analyses

Tests for significance of correlations between different variables (including proportions of schizonts positive for MSPDBL2, and individual gene FPKM values), or estimations of odds ratios and significance of associations between categorical variables, were conducted using a combination of R, Epi-Info and Prism software. Differential gene expression analysis was carried out in DESeq2 (24), focusing on the distributions of derived FPKM values for each gene as defined above.

## Results

### Wide variation in proportions of schizonts expressing MSPDBL2 in clinical isolates

MSPDBL2 protein expression was examined in mature schizonts (each with at least eight nuclei) developed in the first cultured *ex vivo* cycle of each of 96 clinical isolates sampled from malaria patients in four different endemic countries in West Africa. In most isolates, less than 1% of all mature schizonts were positive for MSPDBL2 (overall median of 0.6%), but the frequency distribution was highly skewed as some had much higher proportions (Figure 1 and Supplementary Table S2). Nine isolates had more than 3% schizonts positive, including one that had 73% positive. There were no significant differences in the distributions among different countries (Kruskal-Wallis test and pairwise Mann-Whitney tests non-significant), although the two isolates with highest proportions were from Senegal (Figure 1). The overall distribution was compared to that previously reported for a panel of 12 long-term laboratory-adapted *P. falciparum* lines originally isolated from more diverse sources (6), and this was not significantly different (Mann-Whitney test, P = 0.52). This shows that a wide range of MSPDBL2-positive schizonts, with most isolates having very low proportions positive, is a natural feature of expression in endemic *P. falciparum* populations rather than one which has been selected by laboratory culture.

**Figure 1.**
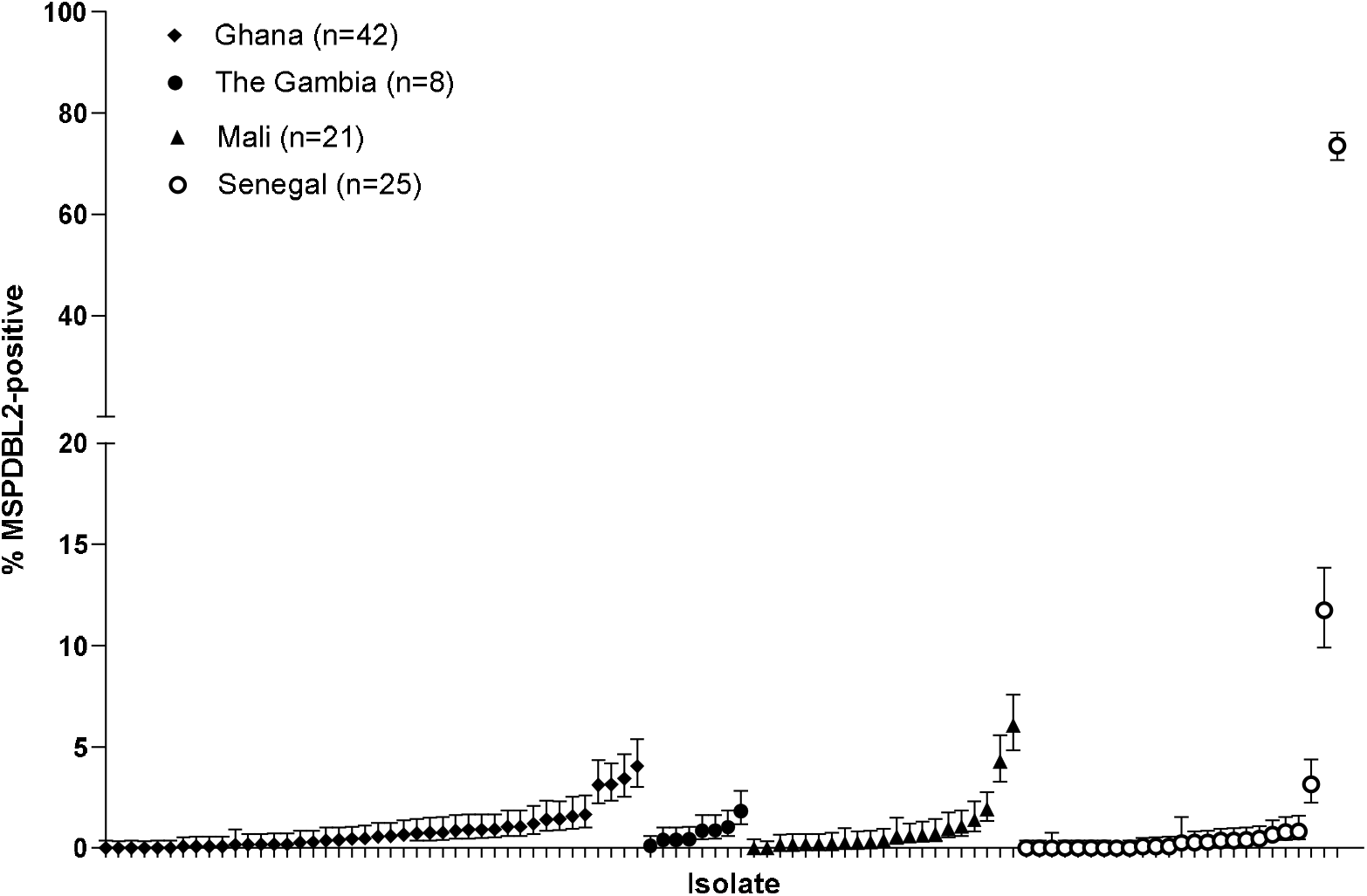
Variable proportions of *P. falciparum* schizonts expressing MSPDBL2 in ex *vivo* cultures of 96 clinical isolates from four West African countries. Samples were from malaria patients in Ghana n=42, The Gambia n=8, Senegal n=25, and Mali n=21. In each isolate, approximately 1000 mature schizonts containing eight or more nuclei were scored in immunofluorescence assays using polyclonal mouse serum against a conserved N-terminal region of MSPDBL2. Eighteen of the isolates had no schizonts positive, and the median across all isolates was only 0.6%, but proportions were highly skewed and nine isolates had more than 3% of schizonts positive. Although the two isolates with highest proportions were from Senegal, there were no significant differences in the overall distributions among different countries by non-parametric rank sum tests.

**Figure 2.**
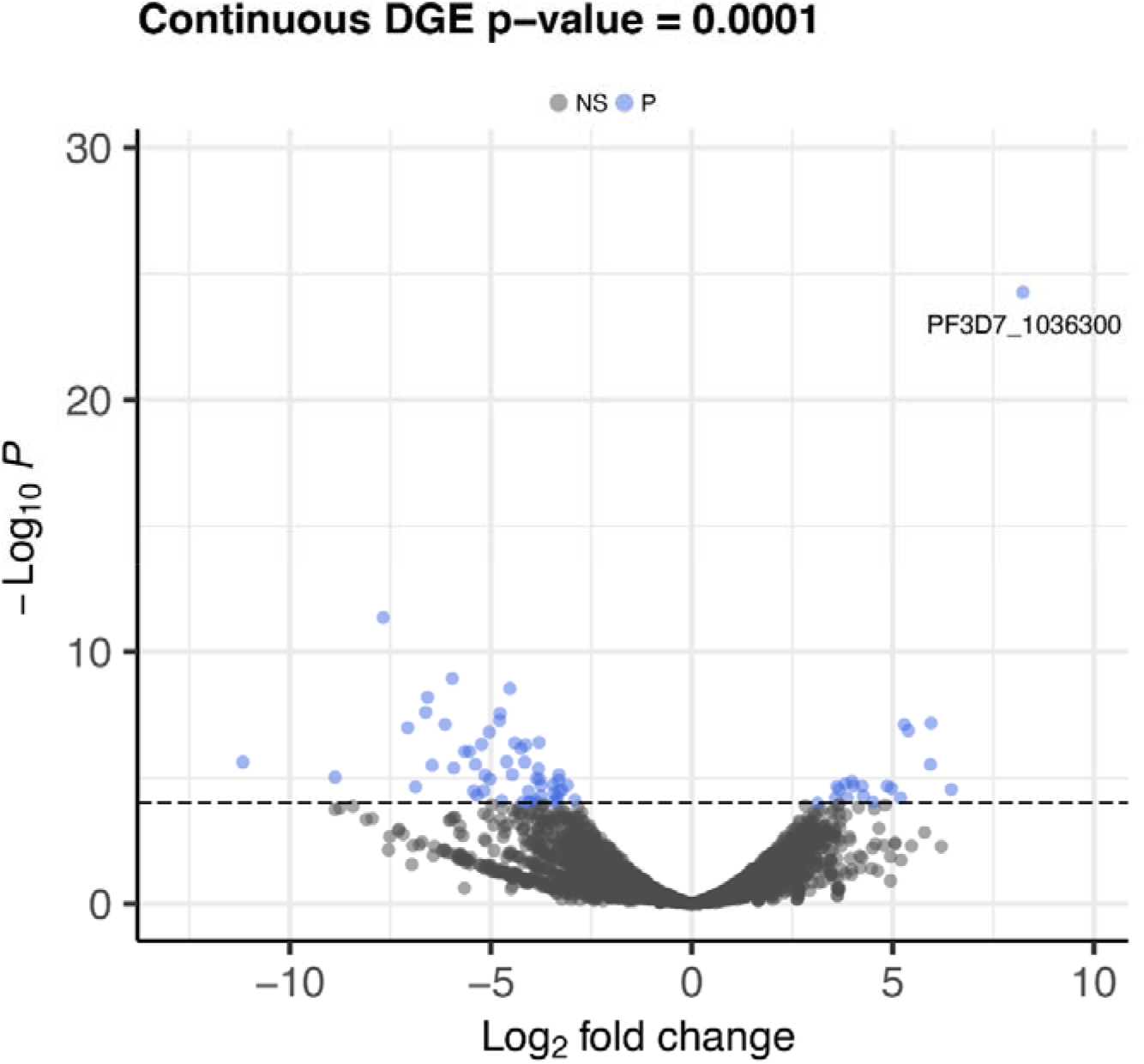
Transcriptome analysis identifies *P. falciparum* genes showing correlation with MSPDBL2 expression in *ex vivo* cultured clinical isolates. RNA-seq was performed on schizont-enriched cultures of 17 West African clinical isolate in the first ex *vivo* cycle, and across the isolates FPKM transcript levels of each gene were tested for correlation with the proportion of schizonts expressing MSPDBL2. Blue shading indicates those that have the most significant differential gene expression (DGE, P < 10^−4^), and genes positively correlated at this level of significance are listed in Table 1. A broader set of genes correlated positively at a slightly lower level of individual significance (P < 0.001, Supplementary Table S4), and those that are negatively correlated at that level of significance are also listed separately (Supplementary Table S5).

### Transcriptomes of schizont-enriched ex *vivo* cultures with a wide range in proportions of MSPDBL2-positive schizonts

To explore whether particular parasite gene transcripts are associated with the proportions of MSPDBL2-positive frequencies, whole transcriptome RNA-seq analysis was performed for 17 of the clinical isolates that had sufficient schizont-enriched material, representing a wide range of MSPDBL2 expression (zero to 73% of schizonts positive, Supplementary Table S1). Sequencing of the cDNA libraries yielded a mean of 2.1 × 10^6^ Illumina short reads for each isolate, and most of these reads aligned to the 3D7 reference genome sequence (Supplementary Table S3), enabling read depth analysis of relative expression of individual genes after normalisation for the total number of reads for each isolate (Supplementary Figure S1). The FPKM values across all genes were first compared to published RNA-seq data from tightly synchronised *P. falciparum* 3D7 parasites sampled at seven timepoints post invasion (0, 8, 16, 24, 32, 40, and 48 hours, the last of which may include some next cycle reinvasion) (27), confirming that Spearman’s rank correlations expression profiles in all samples had strongest correlations with schizont stage parasites (13 isolates correlated most strongly with the 40-hour timepoint and four isolates with the 32-hour timepoint, Supplementary Figure S1).

### Gene transcripts correlating with variable proportions of MSPDBL2-positive schizonts among clinical isolates

Among the 17 isolates with RNA-seq data, individual gene FPKM relative expression values were tested for correlation with MSPDBL2 IFA expression. To scan for significantly correlated genes, a P value cut off of <0.001 was used which identified 52 genes with increased expression (including the *mspdbl2* gene which was by far the most positively correlated as expected, Supplementary Table S4), and 130 genes with negatively correlated expression (Supplementary Table S5). Aside from *mspdbl2* itself, 12 (24%) of the other 51 genes with higher expression were previously listed as having known or suspected roles in gametocytogenesis, whereas of the 130 genes with lower expression, only 10 (7%) were listed with known or suspected gametocytogenesis involvement. An odds ratio of 3.7 (95% CI, 1.5 - 9.2, P = 0.0054) on these proportions indicates a significant skew in gametocytogenesis-related genes being more likely to be positively rather than negatively correlated with proportions of MSPDBL2-positive schizonts.

Using a more stringent correlation value cut off of P <10^−4^ to focus on genes having most highly significant correlations with proportions of schizonts expressing MSPDBL2, 19 genes are identified as positively correlated, of which 9 (47%) were previously indicated as having known or suspected roles in gametocytogenesis (Table 1). At this level of correlation significance, 51 genes are negatively correlated (Supplementary Table S3), of which only one (2%) was previously indicated as gametocytogenesis-related. This indicates a very highly significant skew in gametocytogenesis-related genes being positively rather than negatively correlated with proportions of MSPDBL2-positive schizonts, yielding an odds ratio of 45.0 (95% CI, 5.1 – 396.0, P = 1.2 × 10^−5^).

**Table 1.** *P. falciparum* genes with most highly significantly increased expression (P < 10 ^4^) correlating with proportions of MSPDBL2-positive schizonts in clinical isolates.

| Gene ID | P-value | Gene product description |
| --- | --- | --- |
| *PF3D7_1036300* | $5.4 \times 10^{-25}$ | *duffy binding-like merozoite surface protein 2 (mspdbl2)* |
| **PF3D7_1476600** | $6.7 \times 10^{-08}$ | *Plasmodium* exported protein |
| PF3D7_1474000 | $7.6 \times 10^{-08}$ | conserved *Plasmodium* protein |
| **PF3D7_1102500** | $1.3 \times 10^{-07}$ | *gexp02*, *Plasmodium* exported protein (PHISTb) |
| PF3D7_1461800 | $3.0 \times 10^{-06}$ | conserved *Plasmodium* protein |
| PF3D7_1445700 | $1.4 \times 10^{-05}$ | conserved *Plasmodium* protein |
| PF3D7_0814200 | $1.7 \times 10^{-05}$ | DNA/RNA-binding protein Alba 1 |
| **PF3D7_1466200** | $2.0 \times 10^{-05}$ | early gametocyte enriched phosphoprotein EGXP |
| **PF3D7_0114000** | $2.1 \times 10^{-05}$ | GEXP06, exported protein family 1 |
| **PF3D7_1372100** | $2.1 \times 10^{-05}$ | *Plasmodium* exported protein (PHISTb) |
| **PF3D7_0215000** | $2.2 \times 10^{-05}$ | acyl-CoA synthetase |
| PF3D7_1362700 | $2.6 \times 10^{-05}$ | conserved *Plasmodium* protein |
| **PF3D7_1473700** | $2.8 \times 10^{-05}$ | nucleoporin NUP116/NSP116, putative |
| PF3D7_0829400 | $2.8 \times 10^{-05}$ | prolyl 4-hydroxylase subunit alpha, putative |
| PF3D7_1027300 | $5.2 \times 10^{-05}$ | Peroxiredoxin, nuclear protein |
| PF3D7_0515000 | $6.2 \times 10^{-05}$ | pre-mRNA-splicing factor CWC2, putative |
| **PF3D7_1346800** | $6.2 \times 10^{-05}$ | Pfs47, 6-cysteine protein |
| PF3D7_1132600 | $6.4 \times 10^{-05}$ | pre-mRNA-splicing factor 38A, putative |
| **PF3D7_1477700** | $9.1 \times 10^{-05}$ | *Pfg14.748*, *Plasmodium* exported protein (PHISTa) |
| PF3D7_1431400 | $9.9 \times 10^{-05}$ | surface-related antigen SRA |

Independently, a previous study has investigated the transcriptomic profiles of cultured schizonts of the transgenic *P. falciparum* parasite line 164/TdTom, comparing preparations enriched for sexually-committed versus asexually-committed schizonts (28). The data from this study on the PlasmoDB genomics resource site (29) shows relative transcript levels accessible for all except one of the 19 genes that were most highly positively correlated with MSPDBL2 expression in the present study. Fifteen (83%) of these 18 genes had higher expression in the sexually-committed schizont preparation compared to the asexual schizont preparation, a significantly positive skew compared to random expectations (P < 0.05).

### Genes expressed in correlation with *mspdbl2* gene expression in schizont-enriched cultures of clinical isolates

To complement the above scan based on proportions of MSPDBL2-positive schizonts, the varying transcript levels of *mspdbl2* (FPKM values) among the clinical isolates were analysed to scan for other genes with correlated expression. Using a correlation significance value of P < 0.001 as cut-off identified 41 genes with positively correlated expression (Supplementary Table S6), many of which were also correlated with proportions of MSPDBL2-positive schizonts (16 correlated at P < 0.001, 30 correlated at P < 0.01), and 31 genes had negatively correlated expression (Supplementary Table S7).

Of the 41 genes positively correlated with *mspdbl2* transcript expression, 11 (27%) were previously identified as potentially gametocytogenesis-related, in comparison to only 3 (10%) of the 31 negatively correlated genes, giving an odds ratio of 3.4 (95% CI, 0.9 – 13.6, P = 0.06). Using a higher level of cut-off for correlation significance (P <10^−4^), 19 genes were positively correlated with transcript levels of *mspdbl2* (Table 2), of which 8 (42%) were previously identified as potentially gametocytogenesis-related, in comparison to 2 (15%) of 13 genes negatively associated with *mspdbl2,* giving an odds ratio of 4.0 (95% CI, 0.7 – 23.3, P = 0.11). In summary, the analysis based on *mspdbl2* transcript levels gives broadly similar results to the analysis based on MSPDBL2 positive schizont proportions, but the excess proportions of positively correlated gametocyte-related genes are less significant.

**Table 2.** *P. falciparum* genes with most significantly increased expression (P < 10 ^−4^) correlating with *mspdbl2* transcript levels measured by FPKM in transcriptomes of clinical isolates.

| Gene ID | P-value | Gene product description |
| --- | --- | --- |
| **PF3D7_0114000\*** | $2.6 \times 10^{-9}$ | GEXP06, exported protein family 1 |
| PF3D7_1362700* | $2.6 \times 10^{-8}$ | conserved *Plasmodium* protein |
| **PF3D7_1466200\*** | $2.2 \times 10^{-7}$ | early gametocyte enriched phosphoprotein EGXP |
| **PF3D7_1472200** | $3.1 \times 10^{-7}$ | histone deacetylase, putative |
| **PF3D7_1467600\*** | $6.0 \times 10^{-7}$ | conserved *Plasmodium* protein |
| PF3D7_0214300* | $1.1 \times 10^{-6}$ | conserved *Plasmodium* protein |
| PF3D7_1027300* | $2.3 \times 10^{-6}$ | peroxiredoxin |
| PF3D7_1461800* | $2.5 \times 10^{-6}$ | conserved *Plasmodium* protein |
| **PF3D7_1473700\*** | $2.9 \times 10^{-6}$ | nucleoporin NUP116/NSP116, put |
| PF3D7_1361200* | $8.4 \times 10^{-6}$ | conserved *Plasmodium* protein |
| PF3D7_1474000* | $1.0 \times 10^{-5}$ | conserved *Plasmodium* protein |
| PF3D7_0501400 | $1.5 \times 10^{-5}$ | interspersed repeat antigen |
| **PF3D7_0801900** | $2.0 \times 10^{-5}$ | lysine-specific histone demethylase, put |
| **PF3D7_1408200** | $4.9 \times 10^{-5}$ | AP2 domain transcription factor AP2-G2 |
| PF3D7_0207800 | $5.3 \times 10^{-5}$ | serine repeat antigen 3 |
| PF3D7_1235300 | $7.1 \times 10^{-5}$ | CCR4-NOT transcription complex s4, putative |
| PF3D7_0519500 | $7.4 \times 10^{-5}$ | CCR4 domain-containing protein 1, putative |
| PF3D7_1228300 | $7.5 \times 10^{-5}$ | NIMA related kinase 1 |
| **PF3D7_1134600** | $8.7 \times 10^{-5}$ | zinc finger protein, putative |

## Discussion

This study shows that *P. falciparum* in diverse clinical isolates have a wide range of MSPDBL2 expression positivity in mature schizonts in the first *ex vivo* cycle of development, with most isolates having very low proportions positive. The frequency distribution is similar to that previously described for a smaller number of *P. falciparum* laboratory-adapted lines that had been cultured for many years (6), indicating that this is not a result of selection by culture adaptation. Furthermore, the frequency distribution was similar in isolates from each of the four populations sampled here, which have different levels of malaria infection endemicity within West Africa (20, 30), indicating that parasite populations maintain the wide range of MSPDBL2 expression variation within different endemic environments.

Results of the RNA-seq analyses here indicate that MSPDBL2 protein and gene expression in schizonts in the first cycle of development from clinical isolates is significantly correlated with the expression of other particular genes within the *ex vivo* cultures. Particularly, many of the most strongly correlated genes were previously implicated as having known or suspected involvement in the process of gametocytogenesis. This is consistent with expectations from a functional study on effects of the gametocyte development gene gdv1 in assays of an engineered parasite line with highly induced expression of GDV1 (14), which showed significantly increased transcription of *ap2g* as expected, and also *mspdbl2*, as well as a PHISTa gene (PF3D7_1477700). A separate study indicated that levels of the protein encoded by *pfg14_748* increase as parasites develop along the gametocytogenesis pathway, being detectable alongside the early gametocytogenesis marker Pfs16 in parasite cultures before gametocytes were observed to develop (31). In the present study, *pfg14_748* had expression strongly correlated with *mspdbl2*, but although it was previously shown to be induced by GDV1 (14) it is apparently not dependent on expression of AP2-G (25). Several other genes which correlated with *mspdbl2* expression in the present study (including the nucleoprotein gene *nup116*, and the early gametocyte development marker *gexp02)* have been identified as being upregulated by *ap2-g* (25), but *mspdbl2* was not itself identified to be upregulated by *ap2-g*.

Genes correlating with MSPDBL2 expression in clinical isolates include members of the GEXP family encoding proteins involved in protein export occurring during gametocytogenesis (26), particularly *gexp02* (PF3D7_1102500) and *gexp04* (PF3D7_1372100). As well as *gexp02* being known to result from induced *gdv1* expression (14), it has also been shown that *gexp02* is de-repressed in parasites which have conditional knock out of heterochromatin protein 1 (HP1), presumably due to the resulting activation of *gdv1* (32). However, the correlating expression with *gexp02* and other gametocytogenesis genes here does not indicate that *mspdbl2* is active in the process of sexual commitment or that it is a specific marker. It should be noted that relevant analysis of *gdv1* transcript levels are not possible from the double-stranded cDNA transcriptome data in the present study, as *gdv1* is usually repressed by abundant antisense transcripts initiated from the 3’-intergenic region (14, 33, 34), and ratios of sense to antisense transcripts could only be determined by strand-specific sequencing.

Clearly, if *mspdbl2* is directly involved in gametocytogenesis, it would be expected that *ap2-g* would be identified as differentially expressed in isolates with higher *mspdbl2* expression. As this was not the case in this study, it is alternatively possible that the expression of *mspdbl2* occurs in some parasites in parallel to the sexual commitment process. It is important to note that the correlation with some involved in gametocytogenesis in the present study is at the bulk transcriptome level, so that *mspdbl2* expression might not be within the same individual parasites, but ones that tend to occur in the same bulk population. The only published single cell transcriptome data on *P. falciparum* schizonts are from a laboratory clone in which hardly any schizonts express MSPDBL2, so are not informative on co-expression between the *mspdbl2* gene and others (6, 35). Future insights from single cell transcriptome data will require analysis of *P. falciparum* isolates that have a substantial proportion of schizonts expressing MSPDBL2.

Functional studies would be required to determine whether MSPDBL2 is in any way associated with the sexual commitment process, or whether it is only correlated at the population level within infections or within cultures due to being co-incidentally upregulated by *gdv1* (14). As MSPDBL2 is a merozoite surface protein expressed in late schizonts, the current study analysed transcriptomes of schizont-enriched preparations from *ex vivo* culture, as focused on in few other studies of *P. falciparum* clinical isolates (10, 36). Other studies have analysed parasite transcriptomes from the earlier stages of intra-erythrocytic development that are present in peripheral circulation (37–40), and a temporal analysis of parasite development through to mature schizonts may be needed to identify gene products with which MSPDBL2 is functionally linked, and resolve whether it is a marker of an important parasite subpopulation that could be targeted for vaccination.

## Acknowledgements

We are grateful to the malaria patients and clinical staff for participation in the study. The sample collection was facilitated by staff at Kintampo Health Research Centre in Ghana, at the National Institute for Public Health in Guinea, at Nioro du Sahel Health Centre in Mali and at Pikine Health Centre in Senegal. We appreciate the support of colleagues at the Medical Research Council Unit in The Gambia, the Laboratory of Bacteriology and Virology, Le Dantec Hospital in Senegal, The University of Bamako in Mali, Noguchi Memorial Institute for Medical Research and the University of Ghana, and the London School of Hygiene and Tropical Medicine in enabling this work. This study was supported by an ERC Advanced Award (grant AdG-2011-294428), a Leverhulme-Royal Society Africa Award (grant AA110050), a BBSRC PhD studentship within the London Interdisciplinary Doctoral Training Programme (www.lido-dtp.ac.uk), and an MRC project grant (MR/S009760/1).

