## Supplementary material for "Transcriptome analysis of diverse *Plasmodium falciparum* clinical isolates identifies genes correlating with highly variable expression of merozoite surface protein MSPDBL2": Table S3

**Supplementary Table S3.** MSPDBL2 protein expression and RNA-seq details of 17 clinical isolates.

| **Sample ID** | **MSPDBL2 IFA positive** | **RIN** | **Library concentration (nM)** | **Total reads (x10^6^)** | **Aligning to *Pf* reference genome (%)** | **GEO accession number** |
| --- | --- | --- | --- | --- | --- | --- |
| INV236 | 73.0% | N/A | N/A | 0.8 | 82.0 | GSM5897922 |
| INV374 | 0.4% | 9.8 | 13.1 | 1.1 | 84.9 | GSM5897923 |
| INV395 | 6.5% | 7.6 | 7.80 | 3.0 | 84.9 | GSM5897924 |
| INV396 | 2.0% | 8.2 | 8.60 | 3.8 | 88.2 | GSM5897925 |
| INV401 | 1.1% | 7.9 | 8.10 | 2.0 | 84.6 | GSM5897926 |
| INV406 | 0.6% | 8.8 | 12.8 | 1.6 | 87.5 | GSM5897927 |
| INV410 | 1.0% | 8.1 | 10.8 | 2.6 | 83.9 | GSM5897928 |
| INV419 | 0.0% | 7.2 | 15.3 | 2.4 | 80.3 | GSM5897929 |
| INV425 | 3.7% | 8.2 | 20.1 | 1.4 | 84.8 | GSM5897930 |
| INV440 | 0.7% | 7.4 | 11.7 | 2.1 | 84.6 | GSM5897931 |
| INV445 | 0.2% | 6.4 | 14.0 | 1.8 | 83.3 | GSM5897932 |
| INV446 | 0.3% | 6.5 | 11.6 | 2.0 | 85.4 | GSM5897933 |
| INV448 | 0.0% | 9.0 | 15.5 | 2.9 | 77.6 | GSM5897934 |
| INV449 | 0.8% | 8.7 | 12.5 | 3.2 | 83.6 | GSM5897935 |
| INV450 | 0.1% | 6.7 | 18.2 | 1.7 | 82.8 | GSM5897936 |
| INV455 | 0.2% | 6.0 | 13.7 | 1.4 | 86.9 | GSM5897937 |
| INV459 | 0.0% | 8.2 | 10.1 | 2.3 | 82.5 | GSM5897938 |
