## Supplementary material for "Transcriptome analysis of diverse *Plasmodium falciparum* clinical isolates identifies genes correlating with highly variable expression of merozoite surface protein MSPDBL2": Table S4

**Supplementary Table S4. List of genes with expression in clinical isolates positively correlating with proportions of schizonts expressing MSPDBL2 protein by IFA at significance values of P < 0.001**

| **Gene ID** | **P-value** | **Gene description** |
| --- | --- | --- |
| *PF3D7_1036300* | *5.4 E-25* | *duffy binding-like merozoite surface protein 2* |
| **PF3D7_1476600** | 6.7 E-08 | *Plasmodium* exported protein UF |
| PF3D7_1474000 | 7.6 E-08 | conserved *Plasmodium* protein UF |
| **PF3D7_1102500** | 1.3 E-07 | *Plasmodium* exported protein (PHISTb) UF |
| PF3D7_1461800 | 3.0 E-06 | conserved *Plasmodium* protein UF |
| PF3D7_1445700 | 1.4 E-05 | conserved *Plasmodium* protein UF |
| PF3D7_0814200 | 1.7 E-05 | DNA/RNA-binding protein Alba 1 |
| **PF3D7_1466200** | 2.0 E-05 | early gametocyte enriched phosphoprotein EGXP |
| **PF3D7_0114000** | 2.1 E-05 | exported protein family 1 |
| **PF3D7_1372100** | 2.1 E-05 | *Plasmodium* exported protein (PHISTb) UF |
| **PF3D7_0215000** | 2.2 E-05 | acyl-CoA synthetase |
| PF3D7_1362700 | 2.6 E-05 | conserved *Plasmodium* protein UF |
| **PF3D7_1473700** | 2.8 E-05 | nucleoporin NUP116/NSP116, put |
| PF3D7_0829400 | 2.8 E-05 | prolyl 4-hydroxylase subunit alpha, put |
| PF3D7_1027300 | 5.2 E-05 | peroxiredoxin |
| PF3D7_0515000 | 6.2 E-05 | pre-mRNA-splicing factor CWC2, put |
| **PF3D7_1346800** | 6.2 E-05 | 6-cysteine protein |
| PF3D7_1132600 | 6.4 E-05 | pre-mRNA-splicing factor 38A, put |
| **PF3D7_1477700** | 9.1 E-05 | *Plasmodium* exported protein (PHISTa) UF |
| PF3D7_1431400 | 1.0 E-04 | surface-related antigen SRA |
| PF3D7_1140200 | 1.1 E-04 | conserved *Plasmodium* protein UF |
| **PF3D7_1467600** | 1.2 E-04 | conserved *Plasmodium* protein UF |
| PF3D7_1130200 | 1.2 E-04 | 60S ribosomal protein P0 |
| PF3D7_0202000 | 1.3 E-04 | knob-associated histidine-rich protein |
| PF3D7_1429100 | 1.4 E-04 | apicoplast ribosomal protein L15 precursor, put |
| **PF3D7_0315600** | 1.7 E-04 | zinc finger protein, put |
| PF3D7_0611600 | 1.8 E-04 | basal complex transmembrane protein 1 |
| PF3D7_0927300 | 1.9 E-04 | fumarate hydratase |
| PF3D7_0815500 | 1.9 E-04 | conserved *Plasmodium* protein UF |
| PF3D7_1236200 | 2.3 E-04 | conserved *Plasmodium* protein UF |
| PF3D7_0403600 | 2.3 E-04 | conserved *Plasmodium* protein UF |
| PF3D7_1317000 | 2.7 E-04 | U4/U6.U5 tri-snRNP-associated protein 2 put |
| PF3D7_1233200 | 2.9 E-04 | conserved *Plasmodium* protein UF |
| PF3D7_1437900 | 3.0 E-04 | HSP40, subfamily A |
| PF3D7_0419600 | 3.5 E-04 | ran-specific GTPase-activating protein 1, put |
| PF3D7_0912900 | 4.9 E-04 | 26S proteasome regulatory subunit RPN8, put |
| PF3D7_1432800 | 4.9 E-04 | HP12 protein homolog, put |
| PF3D7_0208000 | 5.1 E-04 | serine repeat antigen 1 |
| PF3D7_1361200 | 5.3 E-04 | conserved *Plasmodium* protein UF |
| PF3D7_1136300 | 5.4 E-04 | tudor staphylococcal nuclease |
| PF3D7_1312100 | 5.7 E-04 | GYF domain-containing protein, put |
| PF3D7_0318900 | 5.9 E-04 | conserved protein UF |
| PF3D7_0527500 | 6.2 E-04 | Hsc70-interacting protein |
| PF3D7_1428300 | 6.3 E-04 | proliferation-associated protein 2g4, put |
| PF3D7_1002400 | 6.4 E-04 | transformer-2 protein homolog beta, put |
| PF3D7_0420100 | 7.6 E-04 | serine/threonine protein kinase RIO2 |
| **PF3D7_0406200** | 8.6 E-04 | sexual stage-specific protein precursor *pfs16* |
| PF3D7_1134000 | 8.9 E-04 | heat shock protein 70 |
| PF3D7_0106900 | 9.0 E-04 | cytidylyltransferase, put |
| PF3D7_0408500 | 9.2 E-04 | NYN domain-containing protein, put |
| PF3D7_1145400 | 9.3 E-04 | dynamin-like protein |
| PF3D7_0214300 | 9.94E-04 | conserved *Plasmodium* protein UF |

Genes identified with expression that correlated with MSPDBL2 expression treated as a continuous variable. Using the p-value cut-off of <0.001, 52 genes were identified to be highly significantly positively correlated (*mspdbl2* itself is the most highly significant, as expected). Of the 51 genes aside from *mspdbl2*, 12 (24%) were previously identified as potentially gametocytogenesis-associated (gene ID highlighted in bold)(Supplementary Table S1), compared to only 10 (7%) of genes among the 130 showing lower expression with increasing MSPDBL2 protein expression. UF: protein has unknown function. Put: putative.
