## Supplementary material for "Transcriptome analysis of diverse *Plasmodium falciparum* clinical isolates identifies genes correlating with highly variable expression of merozoite surface protein MSPDBL2": Table S5

**Supplementary Table S5. List of genes with expression in clinical isolates negatively correlating with proportions of schizonts expressing MSPDBL2 protein by IFA at significance values of P < 0.001**

| **Gene ID** | **P-value** | **Gene product Description** |
| --- | --- | --- |
| PF3D7_0805200 | 4.32E-12 | gamete release protein, putative |
| PF3D7_0311500 | 1.13E-09 | conserved protein, unknown function |
| PF3D7_0501600 | 2.87E-09 | rhoptry-associated protein 2 |
| PF3D7_1416500 | 6.29E-09 | NADP-specific glutamate dehydrogenase |
| PF3D7_0202500 | 2.47E-08 | early transcribed membrane protein 2 |
| PF3D7_0610400 | 2.80E-08 | histone H3 |
| PF3D7_1345100 | 7.44E-08 | thioredoxin 2 |
| PF3D7_1477800 | 1.01E-07 | acyl-CoA binding protein |
| PF3D7_1114100 | 1.52E-07 | rhomboid protease ROM1 |
| PF3D7_1028200 | 3.89E-07 | RING zinc finger protein, putative |
| PF3D7_0721900 | 4.05E-07 | V-type ATPase V0 subunit e, putative |
| PF3D7_1312500 | 4.66E-07 | conserved Plasmodium protein, unknown function |
| PF3D7_0825200 | 4.94E-07 | translation initiation factor IF-3 |
| PF3D7_1321400 | 6.64E-07 | palmitoyltransferase DHHC8, putative |
| PF3D7_0112900 | 8.92E-07 | Plasmodium exported protein, unknown function |
| PF3D7_1401400 | 8.93E-07 | early transcribed membrane protein 14.1 |
| PF3D7_1456800 | 2.29E-06 | V-type H(+)-translocating pyrophosphatase, putative |
| PF3D7_1240100 | 2.36E-06 | early transcribed membrane protein 12 |
| PF3D7_0423800 | 2.37E-06 | cysteine-rich protective antigen |
| PF3D7_0611700 | 2.94E-06 | 60S ribosomal protein L39 |
| PF3D7_1455100 | 3.19E-06 | protein tyrosine phosphatase, putative |
| PF3D7_0402100 | 4.05E-06 | Plasmodium exported protein (PHISTb), unknown function |
| PF3D7_0211600 | 4.31E-06 | UDP-N-acetylglucosamine transferase subunit ALG14, putative |
| PF3D7_1302300 | 7.37E-06 | Plasmodium exported protein, unknown function |
| PF3D7_1339400 | 7.56E-06 | cytochrome c oxidase subunit ApiCOX14, putative |
| PF3D7_0730800 | 7.83E-06 | Plasmodium exported protein, unknown function |
| PF3D7_1102700 | 9.31E-06 | early transcribed membrane protein 11.1 |
| PF3D7_1229400 | 1.06E-05 | macrophage migration inhibitory factor |
| PF3D7_0605900 | 1.12E-05 | long chain polyunsaturated fatty acid elongation enzyme, putative |
| PF3D7_1110000 | 1.13E-05 | conserved Plasmodium protein, unknown function |
| PF3D7_1112000 | 1.24E-05 | conserved protein, unknown function |
| PF3D7_1026500 | 1.83E-05 | conserved Plasmodium protein, unknown function |
| PF3D7_1315600 | 1.92E-05 | CDP-diacylglycerol--inositol 3-phosphatidyltransferase |
| PF3D7_0418500 | 1.95E-05 | trafficking protein particle complex subunit 3, putative |
| PF3D7_0706700 | 2.26E-05 | DNA mismatch repair protein MSH2, putative |
| PF3D7_1147400 | 3.08E-05 | conserved Plasmodium protein, unknown function |
| PF3D7_0606800 | 3.23E-05 | VFT protein |
| PF3D7_1337800 | 3.30E-05 | calcium-dependent protein kinase 5 |
| PF3D7_1102800 | 3.34E-05 | early transcribed membrane protein 11.2 |
| PF3D7_1427700 | 3.42E-05 | conserved Plasmodium protein, unknown function |
| PF3D7_1212600 | 4.04E-05 | SND2 domain-containing protein, putative, unspecified product |
| PF3D7_1345000 | 5.00E-05 | conserved Plasmodium protein, unknown function |
| PF3D7_0817800 | 5.98E-05 | conserved Plasmodium protein, unknown function |
| PF3D7_1238000 | 7.56E-05 | COPI associated protein, putative |
| PF3D7_0507400 | 7.91E-05 | conserved Plasmodium protein, unknown function |
| PF3D7_0919300 | 8.00E-05 | thioredoxin-like protein 1, putative |
| PF3D7_1316200 | 8.40E-05 | ADP-ribosylation factor, putative |
| PF3D7_0805600 | 8.79E-05 | PAP2-like protein, putative |
| PF3D7_1127600 | 9.29E-05 | CRAL/TRIO domain-containing protein, putative |
| PF3D7_1324400 | 9.42E-05 | PRELI domain-containing protein, putative |
| PF3D7_0205500 | 9.56E-05 | DNA-directed RNA polymerase II 16 kDa subunit, putative |
| PF3D7_0501700 | 1.03E-04 | anaphase-promoting complex subunit 3, putative |
| PF3D7_0931200 | 1.11E-04 | selenoprotein |
| PF3D7_1117300 | 1.14E-04 | conserved protein, unknown function |
| PF3D7_1219900 | 1.19E-04 | ribulose-phosphate 3-epimerase, putative |
| PF3D7_1010300 | 1.30E-04 | succinate dehydrogenase subunit 4, putative |
| PF3D7_1243300 | 1.32E-04 | conserved Plasmodium protein, unknown function |
| PF3D7_0932200 | 1.39E-04 | profilin |
| PF3D7_0217800 | 1.47E-04 | 40S ribosomal protein S26 |
| PF3D7_0402900 | 1.54E-04 | probable protein, unknown function |
| PF3D7_1404800 | 1.54E-04 | conserved Plasmodium protein, unknown function |
| PF3D7_0425100 | 1.76E-04 | Plasmodium exported protein (hyp6), unknown function |
| PF3D7_1104700 | 1.95E-04 | DNA-directed RNA polymerase III subunit RPC8, putative |
| PF3D7_0528000 | 2.06E-04 | proteasome maturation factor UMP1, putative |
| PF3D7_0813800 | 2.06E-04 | GDP-mannose 4,6-dehydratase |
| PF3D7_0312300 | 2.24E-04 | 26S proteasome regulatory subunit RPN12, putative |
| PF3D7_1144000 | 2.24E-04 | 40S ribosomal protein S21 |
| PF3D7_1426900 | 2.29E-04 | cytochrome b-c1 complex subunit 6, putative |
| PF3D7_1025800 | 2.37E-04 | conserved protein, unknown function |
| PF3D7_1242900 | 2.46E-04 | mitochondrial import inner membrane translocase subunit TIM13 |
| PF3D7_0702200 | 2.58E-04 | lysophospholipase, putative |
| PF3D7_1467700 | 2.61E-04 | conserved Plasmodium protein, unknown function |
| PF3D7_0721200 | 2.61E-04 | conserved Plasmodium protein, unknown function |
| PF3D7_1478800 | 2.66E-04 | Plasmodium exported protein, unknown function |
| PF3D7_1238900 | 2.73E-04 | protein kinase 2 |
| PF3D7_1433300 | 2.82E-04 | chromatin assembly factor 1 P55 subunit, putative |
| PF3D7_0824700 | 3.01E-04 | lipase maturation factor, putative |
| PF3D7_0301800 | 3.24E-04 | Plasmodium exported protein, unknown function |
| PF3D7_1472600 | 3.24E-04 | protein disulfide-isomerase |
| PF3D7_0613100 | 3.34E-04 | conserved Plasmodium protein, unknown function |
| PF3D7_1132800 | 3.39E-04 | aquaglyceroporin |
| PF3D7_1210900 | 3.43E-04 | GPI mannosyltransferase 1 |
| PF3D7_1366800 | 3.44E-04 | phosphatidylserine synthase, putative |
| PF3D7_0503800 | 3.59E-04 | 60S ribosomal protein L31 |
| PF3D7_1252200 | 3.67E-04 | chitinase |
| PF3D7_0113900 | 3.68E-04 | CX3CL1-binding protein 1 |
| PF3D7_0819900 | 3.68E-04 | U6 snRNA-associated Sm-like protein LSm3, putative |
| PF3D7_0904700 | 3.86E-04 | bacterial histone-like protein |
| PF3D7_1454500 | 3.91E-04 | iron sulfur cluster assembly protein, putative |
| PF3D7_1325000 | 3.94E-04 | U6 snRNA-associated Sm-like protein LSm6, putative |
| PF3D7_1306600 | 3.98E-04 | V-type proton ATPase subunit H, putative |
| PF3D7_1001900 | 4.07E-04 | Plasmodium exported protein (hyp16), unknown function |
| PF3D7_1230400 | 4.09E-04 | ATP-dependent protease subunit ClpQ |
| PF3D7_0320900 | 4.17E-04 | histone H2A.Z |
| PF3D7_0424600 | 4.28E-04 | Plasmodium exported protein (PHISTb), unknown function |
| PF3D7_0219700 | 4.35E-04 | Plasmodium exported protein (PHISTc), unknown function |
| PF3D7_0414300 | 4.37E-04 | Rab5-interacting protein, putative |
| PF3D7_1001500 | 4.41E-04 | early transcribed membrane protein 10.1 |
| PF3D7_1476800 | 4.43E-04 | lysophospholipase, putative |
| PF3D7_0422900 | 4.52E-04 | methyltransferase, putative |
| PF3D7_0916700 | 4.54E-04 | RNA-binding protein musashi, putative |
| PF3D7_0307100 | 4.62E-04 | 40S ribosomal protein S12, putative |
| PF3D7_0422400 | 4.78E-04 | 40S ribosomal protein S19 |
| PF3D7_1200800 | 4.84E-04 | serine/threonine protein kinase, FIKK family |
| PF3D7_0713700 | 4.99E-04 | conserved Plasmodium protein, unknown function |
| PF3D7_1034200 | 5.10E-04 | ribosomal protein L27, putative |
| PF3D7_0617300 | 5.15E-04 | conserved Plasmodium protein, unknown function |
| PF3D7_1010400 | 5.16E-04 | MORN repeat protein, putative |
| PF3D7_0105400 | 5.17E-04 | conserved Plasmodium protein, unknown function |
| PF3D7_1218500 | 5.25E-04 | dynamin-like protein, putative |
| PF3D7_0825400 | 6.17E-04 | ATP synthase-associated protein, putative |
| PF3D7_1102400 | 6.23E-04 | phosphopantothenate--cysteine ligase, putative |
| PF3D7_1133000 | 6.30E-04 | conserved Plasmodium protein, unknown function |
| PF3D7_0204700 | 6.40E-04 | hexose transporter |
| PF3D7_1104100 | 6.59E-04 | syntaxin, Qa-SNARE family |
| PF3D7_1030000 | 6.68E-04 | transcription elongation factor SPT4, putative |
| PF3D7_0202200 | 7.30E-04 | EMP1-trafficking protein |
| PF3D7_0219900 | 7.34E-04 | Plasmodium exported protein, unknown function |
| PF3D7_1332300 | 7.70E-04 | trafficking protein particle complex subunit 2, putative |
| PF3D7_0709000 | 8.13E-04 | chloroquine resistance transporter |
| PF3D7_1201200 | 8.51E-04 | Plasmodium exported protein (PHISTa-like), unknown function |
| PF3D7_1253100 | 8.57E-04 | Plasmodium exported protein (PHISTa), unknown function |
| PF3D7_0103800 | 8.81E-04 | actin-related protein |
| PF3D7_0805400 | 9.13E-04 | N-acetyltransferase, GNAT family, putative |
| PF3D7_0935600 | 9.33E-04 | gametocytogenesis-implicated protein |
| PF3D7_1144100 | 9.39E-04 | mitochondrial large subunit ribosomal protein, putative |
| PF3D7_0711100 | 9.54E-04 | conserved protein, unknown function |
| PF3D7_0506400 | 9.56E-04 | conserved Plasmodium protein, unknown function |
| PF3D7_1105600 | 9.62E-04 | translocon component PTEX88 |
| PF3D7_1331600 | 9.78E-04 | protein tyrosine phosphatase-like protein, putative |

Genes identified with expression that correlated with MSPDBL2 expression treated as a continuous variable. Using the p-value cut-off of <0.001, 130 genes were identified to be highly significantly negatively correlated. UF: protein has unknown function. Put: putative.
