## Supplementary material for "Transcriptome analysis of diverse *Plasmodium falciparum* clinical isolates identifies genes correlating with highly variable expression of merozoite surface protein MSPDBL2": Table S6

**Supplementary Table S6. List of genes with increased expression in clinical isolates positively correlating with *mspdbl2* transcript levels measured by FPKM at significance values of P < 0.001**

| **Gene ID** | **P-value** | | **Product Description** |
| --- | --- | --- | --- |
| **PF3D7_0114000*** | 2.63 E-09 | exported protein family 1 | |
| PF3D7_1362700* | 2.60 E-08 | conserved *Plasmodium* protein, UF | |
| **PF3D7_1466200*** | 2.17 E-07 | early gametocyte enriched phosphoprotein EGXP | |
| **PF3D7_1472200** | 3.09 E-07 | histone deacetylase, putative | |
| **PF3D7_1467600*** | 6.00 E-07 | conserved *Plasmodium* protein, UF | |
| PF3D7_0214300* | 1.11 E-06 | conserved *Plasmodium* protein UF | |
| PF3D7_1027300* | 2.30 E-06 | Peroxiredoxin | |
| PF3D7_1461800* | 2.52 E-06 | conserved *Plasmodium* protein, UF | |
| **PF3D7_1473700*** | 2.89 E-06 | nucleoporin NUP116/NSP116, put | |
| PF3D7_1361200* | 8.35 E-06 | conserved *Plasmodium* protein, UF | |
| PF3D7_1474000* | 1.01 E-05 | conserved *Plasmodium* protein, UF | |
| PF3D7_0501400 | 1.49 E-05 | interspersed repeat antigen | |
| **PF3D7_0801900** | 2.01 E-05 | lysine-specific histone demethylase, put | |
| **PF3D7_1408200** | 4.85 E-05 | AP2 domain transcription factor AP2-G2 | |
| PF3D7_0207800 | 5.33 E-05 | serine repeat antigen 3 | |
| PF3D7_1235300 | 7.10 E-05 | CCR4-NOT transcription complex s4, put | |
| PF3D7_0519500 | 7.43 E-05 | CCR4 domain-containing protein 1, put | |
| PF3D7_1228300 | 7.52 E-05 | NIMA related kinase 1 | |
| **PF3D7_1134600** | 8.65 E-05 | zinc finger protein, putative | |
| **PF3D7_0315600*** | 1.81 E-04 | zinc finger protein, putative | |
| PF3D7_1133700 | 1.27 E-04 | FHA domain-containing protein, put | |
| PF3D7_1236200* | 1.33 E-04 | conserved *Plasmodium* protein, UF | |
| PF3D7_1212700 | 1.49 E-04 | eukaryotic translation initiation factor 3.A, putative | |
| PF3D7_1233200* | 1.57 E-04 | conserved *Plasmodium* protein, UF | |
| PF3D7_1327300 | 2.35 E-04 | conserved *Plasmodium* protein, UF | |
| **PF3D7_1102500*** | 2.40 E-04 | *Plasmodium* exported protein (PHISTb), UF | |
| PF3D7_1469600 | 2.51 E-04 | acetyl-CoA carboxylase | |
| PF3D7_0309200 | 3.31 E-04 | serine/threonine protein kinase, putative | |
| PF3D7_0724100 | 3.94 E-04 | conserved *Plasmodium* protein, UF | |
| PF3D7_0723400 | 4.46 E-04 | conserved *Plasmodium* protein, UF | |
| PF3D7_0829400* | 5.28 E-04 | prolyl 4-hydroxylase subunit alpha, putative | |
| PF3D7_1132400 | 5.86 E-04 | conserved *Plasmodium* membrane protein, UF | |
| PF3D7_1138800 | 6.24 E-04 | WD repeat-containing protein, putative | |
| PF3D7_1142100 | 7.22 E-04 | conserved *Plasmodium* protein, UF | |
| PF3D7_1014300 | 7.43 E-04 | SPRY domain-containing protein, putative | |
| PF3D7_1437200 | 8.05 E-04 | ribonucleoside-diphosphate reductase subunit, | |
| PF3D7_0930300 | 9.74 E-04 | merozoite surface protein 1 | |
| PF3D7_1133800 | 9.79 E-04 | RNA (uracil-5-)methyltransferase, putative | |
| **PF3D7_1148700** | 9.80 E-04 | *Plasmodium* exported protein (PHISTc), UF | |
| PF3D7_0402200 | 9.99 E-04 | surface-associated interspersed protein 4.1 pseudo | |

41 genes identified with expression that correlated with higher *mspdbl2* transcript levels. Genes highlighted bold have known or suspected roles in gametocytogenesis. Asterisks * indicate genes also identified as having higher expression correlating to MSPDBL2 protein expression in schizonts by IFA. UF: protein has unknown function.
