## Supplementary material for "Transcriptome analysis of diverse *Plasmodium falciparum* clinical isolates identifies genes correlating with highly variable expression of merozoite surface protein MSPDBL2": Table S7

**Supplementary Table S7. List of genes with expression in clinical isolates negatively correlating with *mspdbl2* transcript levels measured by FPKM at significance values of P < 0.001**

| Gene ID | P-value | Product Description |
| --- | --- | --- |
| PF3D7_0805200 | 2.04E-08 | gamete release protein, putative |
| PF3D7_1416500 | 4.01E-07 | NADP-specific glutamate dehydrogenase |
| PF3D7_1102700 | 7.05E-07 | early transcribed membrane protein 11.1 |
| PF3D7_0202500 | 2.08E-06 | early transcribed membrane protein 2 |
| PF3D7_0402100 | 1.15E-05 | Plasmodium exported protein (PHISTb), unknown function |
| PF3D7_0605900 | 1.20E-05 | long chain polyunsaturated fatty acid elongation enzyme |
| PF3D7_1401400 | 1.74E-05 | early transcribed membrane protein 14.1 |
| PF3D7_1337800 | 2.15E-05 | calcium-dependent protein kinase 5 |
| PF3D7_0424600 | 2.29E-05 | Plasmodium exported protein (PHISTb), unknown function |
| PF3D7_1238900 | 5.77E-05 | protein kinase 2 |
| PF3D7_1240100 | 7.69E-05 | early transcribed membrane protein 12 |
| PF3D7_1324400 | 8.22E-05 | PRELI domain-containing protein, putative |
| PF3D7_1102800 | 9.96E-05 | early transcribed membrane protein 11.2 |
| PF3D7_0702200 | 1.05E-04 | lysophospholipase, putative |
| PF3D7_1218500 | 1.54E-04 | dynamin-like protein, putative |
| PF3D7_0210000 | 1.81E-04 | secretory complex protein 61 gamma subunit |
| PF3D7_1477800 | 1.96E-04 | acyl-CoA binding protein |
| PF3D7_1401100 | 2.16E-04 | DnaJ protein, putative |
| PF3D7_1476800 | 2.26E-04 | lysophospholipase, putative |
| PF3D7_1404800 | 2.50E-04 | conserved Plasmodium protein, unknown function |
| PF3D7_1102400 | 2.69E-04 | phosphopantothenate--cysteine ligase, putative |
| PF3D7_0916700 | 2.97E-04 | RNA-binding protein musashi, putative |
| PF3D7_0112900 | 3.58E-04 | Plasmodium exported protein, unknown function |
| PF3D7_1472600 | 5.08E-04 | protein disulfide-isomerase |
| PF3D7_0730800 | 5.26E-04 | Plasmodium exported protein, unknown function |
| PF3D7_0206200 | 5.56E-04 | pantothenate transporter |
| PF3D7_1253100 | 5.87E-04 | Plasmodium exported protein (PHISTa), unknown function |
| PF3D7_1245000 | 6.15E-04 | 5-formyltetrahydrofolate cyclo-ligase, putative |
| PF3D7_1312500 | 6.79E-04 | conserved Plasmodium protein, unknown function |
| PF3D7_1104100 | 7.47E-04 | syntaxin, Qa-SNARE family |
| PF3D7_1201200 | 8.95E-04 | Plasmodium exported protein (PHISTa-like) |

31 genes identified with expression that correlated with lower *mspdbl2* transcript levels. UF: protein has unknown function.
