## Supplementary material for "Transcriptome analysis of diverse *Plasmodium falciparum* clinical isolates identifies genes correlating with highly variable expression of merozoite surface protein MSPDBL2": Figure S1

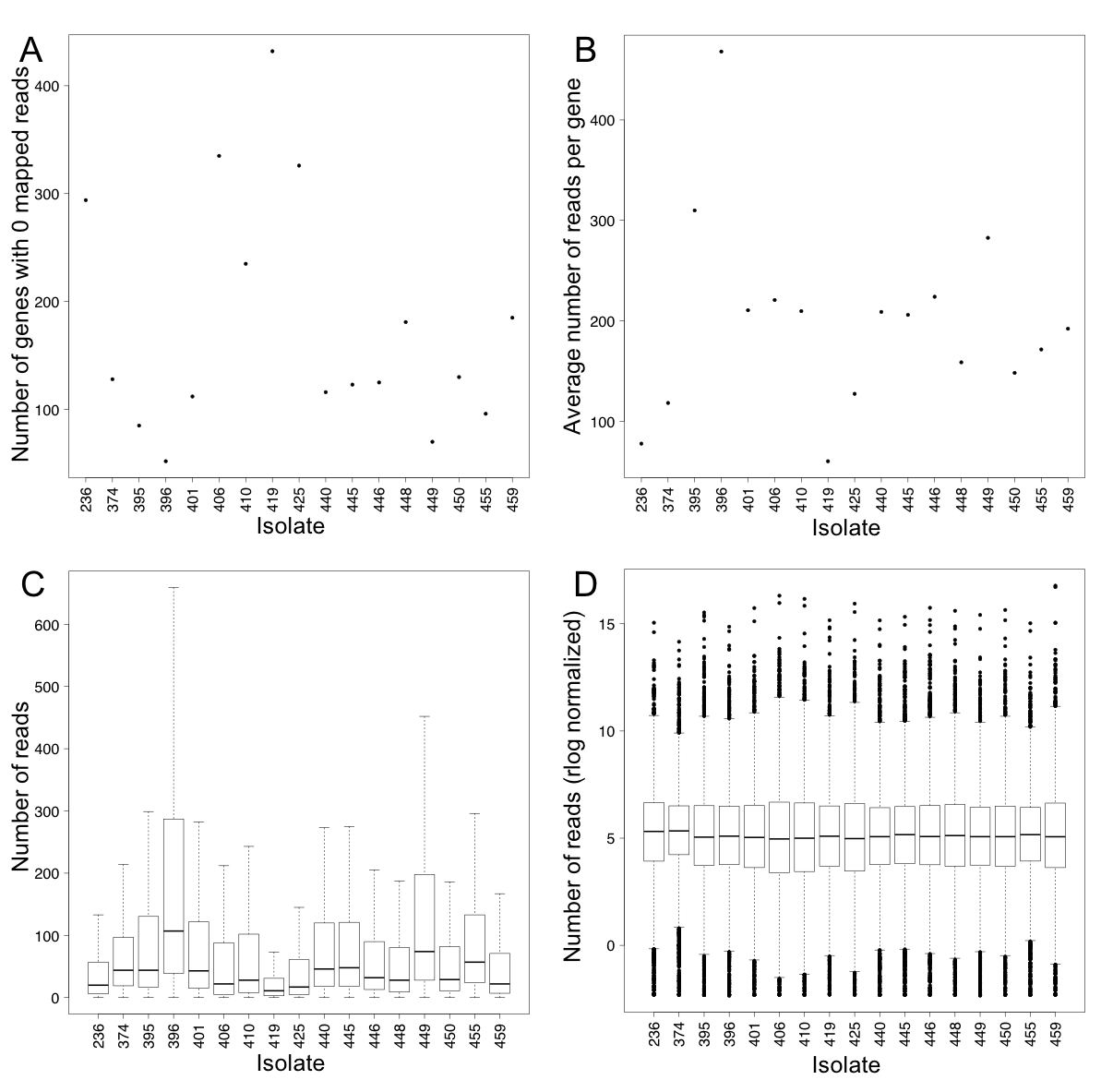

**Supplementary Figure S1. Graphs summarising variation in sequence read depths among individual clinical isolate cDNA libraries and normalisation. A.** The number of genes with zero mapped reads per isolate. **B.** The average number of reads per gene. **C.** Distribution of the numbers of reads per gene based on raw read data. **D.** Variation in number of reads representing genes is normalised by regularised log (rlog) transformation in DESeq2.
