## Supplementary material for "Transcriptome analysis of diverse *Plasmodium falciparum* clinical isolates identifies genes correlating with highly variable expression of merozoite surface protein MSPDBL2": Figure S2

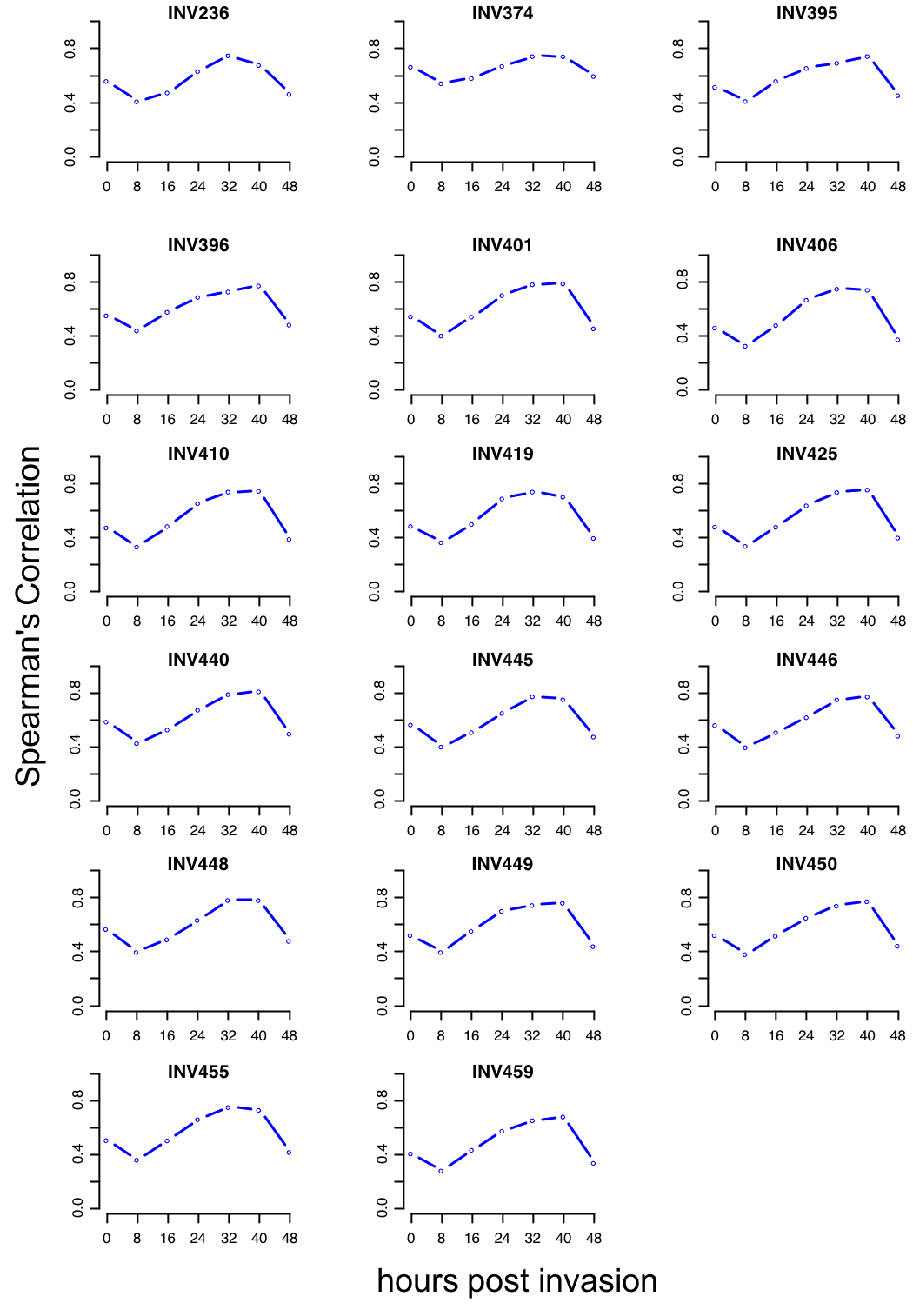

**Supplementary Figure S2.** **Correlations of the RNA-seq profiles of each of the** **17 clinical isolates with different stages of parasite development.** FPKM values for the 17 clinical isolates were correlated with reference data on a laboratory parasite clone across seven timepoints in the *P. falciparum* intraerythrocytic developmental cycle (Otto *et al.* 2010. *Mol. Microbiol.* 76:12-24). All isolate preparations had highest correlation with the reference profile at either 32 or 40 hours post invasion.
